# Muscle weakness precedes atrophy during cancer cachexia and is associated with muscle-specific mitochondrial stress

**DOI:** 10.1101/2021.09.15.460477

**Authors:** Luca J. Delfinis, Catherine A. Bellissimo, Shivam Gandhi, Sara N. DiBenedetto, Megan E. Rosa-Caldwell, Fasih A. Rahman, Michael P. Wiggs, Uwe Schlattner, Joe Quadrilatero, Nicholas P. Greene, Christopher G.R. Perry

**Affiliations:** Muscle Health Research Centre, School of Kinesiology, York University, 4700 Keele Street, Toronto, ON, Canada; Cachexia Research Laboratory, Department of Health, Human Performance and Recreation, University of Arkansas, Fayetteville, USA; Faculty of Applied Health Sciences, Department of Kinesiology, University of Waterloo, Waterloo, Ontario, Canada; Mooney Lab for Exercise, Nutrition, and Biochemistry, Department of Health, Human Performance and Recreation, Baylor University, Waco, TX, USA; Laboratory of Fundamental and Applied Bioenergetics (LBFA) and SFR Environmental and Systems Biology (BEeSy), University of Grenoble Alpes, Grenoble, France

**Keywords:** Cancer cachexia, mitochondria, skeletal muscle

## Abstract

Muscle weakness and wasting are defining features of cancer-induced cachexia. Mitochondrial stress occurs before atrophy in certain muscles, but distinct responses between muscles and across time remains unclear. We aimed to determine the time-dependent and muscle-specific responses to Colon-26 (C26) cancer-induced cachexia in mice. At 2 weeks post-inoculation, the presence of small tumours did not alter body or muscle mass but decreased force production in the quadriceps and diaphragm. Pyruvate-supported mitochondrial respiration was lower in quadriceps while mitochondrial H_2_O_2_ emission was elevated in diaphragm. At 4 weeks, large tumours corresponded to lower body mass, muscle mass, and cross-sectional area of fibers in quadriceps and diaphragm. Force production in quadriceps was unchanged but remained lower in diaphragm vs control. Mitochondrial respiration was increased while H_2_O_2_ emission was unchanged in both muscles vs control. Mitochondrial creatine sensitivity was compromised in quadriceps. These findings indicate muscle weakness precedes atrophy in quadriceps and diaphragm but is linked to heterogeneous mitochondrial alterations. Eventual muscle-specific restorations in force and bioenergetics highlight how the effects of cancer on one muscle do not predict the response in another muscle. Exploring heterogeneous responses of muscles to cancer may reveal new mechanisms underlying distinct sensitivities, or resistance, to cancer cachexia.

## Introduction

Cancer-induced cachexia is a multifactorial syndrome characterized, in part, by a loss of skeletal muscle mass that cannot be fully reversed by conventional nutritional support (1). This condition leads to progressive reductions in functional independence and quality of life (2). Such declines in muscle mass also reduce tolerance to anticancer therapies and overall survivability (3, 4), and is associated with increased hospitalization time (5). 20-80% of cancer patients are thought to develop cachexia depending on the type and stage of cancer (6). However, the time-dependent relationship between muscle atrophy and weakness remains unclear, as does the degree to which this relationship may vary between muscle types. Exploring the natural divergence of muscle responses to cancer may be an opportunistic approach to identify distinct mechanisms underlying muscle weakness and wasting during cancer cachexia.

Contemporary theories posit that muscle wasting during cachexia is induced by circulating factors generated during cancer which trigger protein degradation and loss of myofibrillar proteins through various mechanisms (4, 7, 8). However, recent literature suggests skeletal muscle mitochondria are also subject to damage during cancer cachexia (9–11) and may be direct contributors to either muscle weakness or atrophy. Current literature suggests oxidative phosphorylation is impaired in the soleus, gastrocnemius and plantaris muscle of tumour-bearing mice, while reactive oxygen species (ROS) - in the form of mitochondrial H_2_O_2_ emission (mH_2_O_2_) - can be increased or decreased depending on the muscle and duration of cancer (9, 11, 12). This suggests cellular mechanisms contributing to muscle loss during cancer cachexia may be more complicated than previously believed. Moreover, in the Lewis lung carcinoma (LLC) xenograft mouse model, certain indices of skeletal muscle mitochondrial dysfunction preceded the onset of muscle atrophy, suggesting mitochondria may be a potential therapeutic target (12). This theory was supported by subsequent studies reporting positive effects of the mitochondria-targeting compound SS-31 in preventing certain indices of cachexia in some but not all muscles of the C26 xenograft mouse model (13, 14). However, given the multifactorial contributions to cachexia during cancer, it seems likely the relationship between mitochondria and myopathy may differ between muscle type and throughout cancer progression.

Indeed, skeletal muscle mitochondria are known to be highly adaptable to metabolic stressors and can super-compensate during an energy crisis (15, 16). In this light, the available literature does not provide sufficient information to predict the extent to which cancer will affect individual muscles, particularly in relation to their underlying mitochondrial responses to the systemic stress of this disease. Understanding the time-dependent nature of unique mitochondrial signatures during cancer-induced cachexia might better inform the development of mitochondrial therapies that have so far yielded disparate results across various muscle types in the C26 cancer mouse model (13, 14).

In this study, we compared the time-dependent relationship of muscle dysfunction and mitochondrial bioenergetic responses to cancer between locomotor (quadriceps) and respiratory muscles (diaphragm). In so doing, we employed a careful consideration of mitochondrial substrate titration protocols modeling key parameters governing mitochondrial bioenergetics *in vivo*. Similar assay design considerations have been essential for identifying precise mitochondrial bioenergetic contributions to cellular function in our previous research (17–20). Using the C26 tumour-bearing mouse model, we reveal muscle weakness precedes atrophy in quadriceps and diaphragm. Energetic insufficiencies were more pronounced in quadriceps whereas mitochondrial redox stress was more evident in diaphragm, yet both muscles showed a delayed correction, if not super-compensation, as cancer progressed. These findings demonstrate the effects of cancer on one muscle do not necessarily predict the response in another muscle type. Moreover, the heterogeneous muscle-specific and time-dependent mitochondrial relationships to cancer may represent an opportunity for informing a more targeted approach to developing mitochondrial therapies to improve muscle health in this debilitating disorder.

## Results

### C26 tumour-bearing mice show progressive reductions in body weight and muscle mass

Body weights were reduced 4 weeks after subcutaneous implantations of C26 cells (Figure 1A), while tumour-free body weights progressively decreased beginning at 3 weeks to a net loss of 27% by 4 weeks (Figure 1B, C) at a time of substantial tumour growth (Figure 1D). Tumours grew to ~0.2g at 2 weeks and ~2.2g at 4 weeks (Figure 1E). C26 spleen mass (marker of inflammatory stress) was not different from PBS at 2 weeks but was significantly greater at 4 weeks (Figure 1F). The mass of specific muscles was similar between C26 and PBS at 2 weeks (Figure 1G). At 4 weeks soleus (SOL) mass was similar between C26 and PBS while lower muscle masses were observed in C26 for extensor digitorum longus (EDL; −23%), plantaris (PLA; −20%), tibialis anterior (TA; −26%), gastrocnemius (GA; −21%) and quadriceps (QUAD; −29%) vs PBS (Figure 1H).

**Figure 1.**
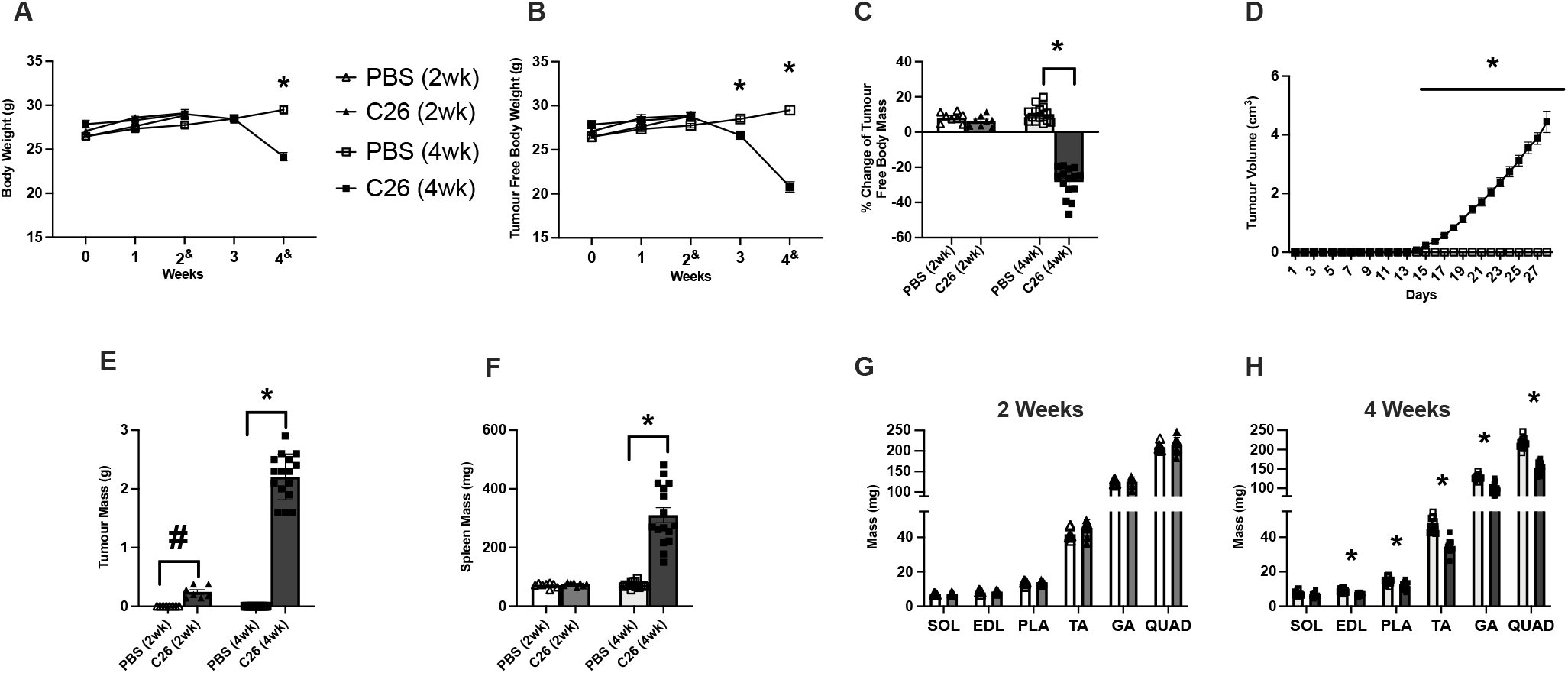
The effects of C26 colon cancer cells implantation on body size, tumour size, muscle mass and force. Analysis of CD2F1 mice with subcutaneous C26 implantations or with PBS were performed. Body weights (*A*, *n*=8-16) and tumour-free body weights (*B*, *n*=8-16) were analyzed every week (2^&^ mice were measured at a 14-17 day window and 4^&^ mice were measured on a 26-29 day window). Percent change in tumour free body weights were analyzed from day 0 to end point (*C*, *n*=8-16). *In vivo* tumour volume measurements were made using calipers (*D*, *n*=16). Tumour mass (*E*, *n*=7-16) and spleen mass (*F*, *n*=8-16) measurements were also completed. Evaluation of hindlimb muscle wet weights were made in the 2-week cohort (*G*, *n*=8) and 4-week cohort (*H*, *n*=16). Results represent mean ± SEM; # *P*<0.05 PBS(2wk) vs C26(2wk); * *P*<0.05 PBS(4wk) vs C26(4wk).

### Force production is reduced prior to atrophy in quadriceps and diaphragm but eventually returns to normal in quadriceps

At 2 weeks, C26 muscle force was lower in quadriceps (Figure 2A) and diaphragm (Figure 2B) relative to PBS as a group main effect. In the quadriceps, there was an interaction whereby C26 at 2 weeks produced less force at 80Hz, 100Hz and 120Hz compared to both PBS groups and the C26 at 4 weeks (not shown). By 4 weeks, quadriceps force in C26 was not different compared to PBS (Figure 2A). In contrast, diaphragm force remained lower relative to PBS control mice at 4 weeks (Figure 2B).

**Figure 2.**
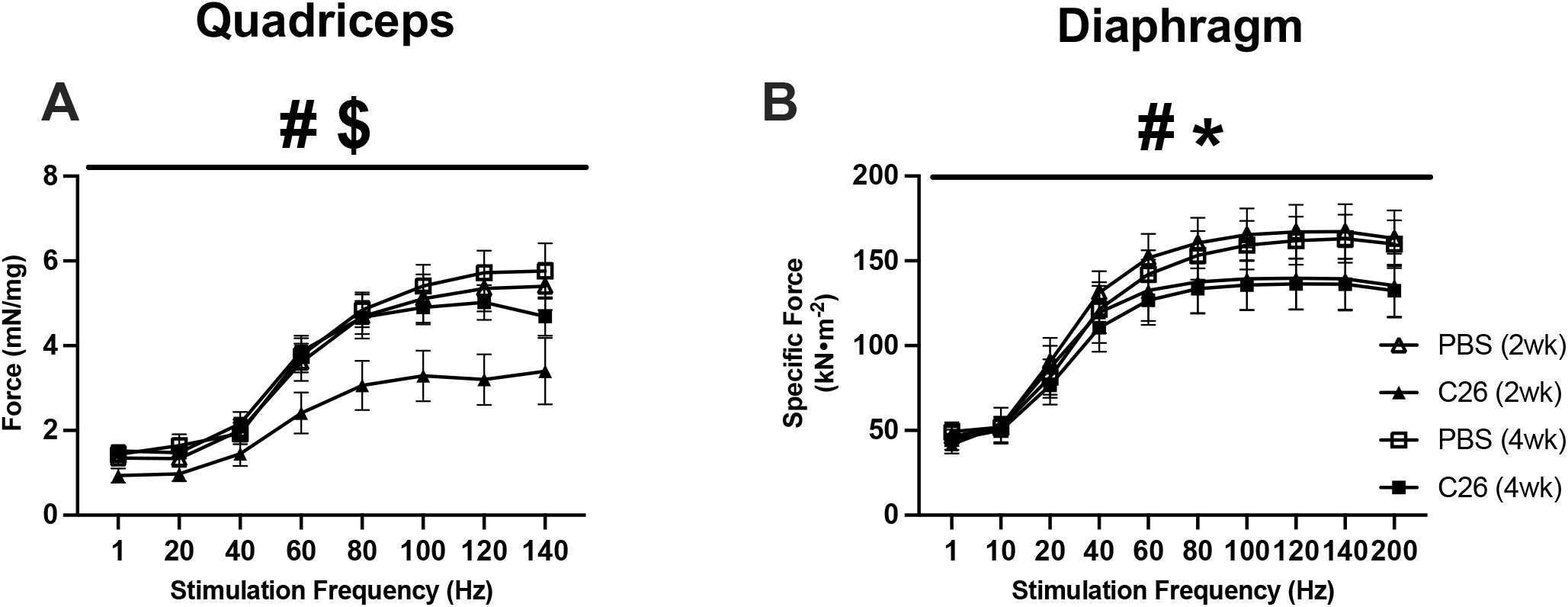
The effects of C26 colon cancer on quadriceps and diaphragm force production. *In situ* quadriceps force production was assessed using the force frequency relationship (*A*, *n*=6-14) and *in vitro* diaphragm force production was also measured using the force frequency relationship (*B*, *n*=6-12). Results represent mean ± SEM; # *P*<0.05 PBS(2wk) vs C26(2wk); * *P*<0.05 PBS(4wk) vs C26(4wk); $ *P*<0.05 C26 (2wk) vs C26 (4wk).

In both quadriceps and diaphragm, fibre CSA was similar between C26 and PBS groups for specific MHC isoforms (Figure 3A-D) and when pooling all MHC isoforms (data not shown) at 2 weeks. However, at 4 weeks, quadriceps muscle exhibited lower CSA in pooled fibres (−40%, *p*<0.05, data not shown) with specific reductions in MHC IIX (−32%) and MHC IIB (−49%) but not the MHC IIA isoform (Figure 3E, F) vs PBS. MHC I-positive fibres were not detected in the quadriceps (Figure 3B, F). At 4 weeks, diaphragm muscle also showed lower CSA in pooled fibres (−31%, *p*<0.05, data not shown) which mirrored changes in MHC I (−28%), MHC IIA (−21%), MHC IIB (−30%) and MHC IIX (−35%) vs PBS at 4 weeks (Figure 3G, H).

**Figure 3.**
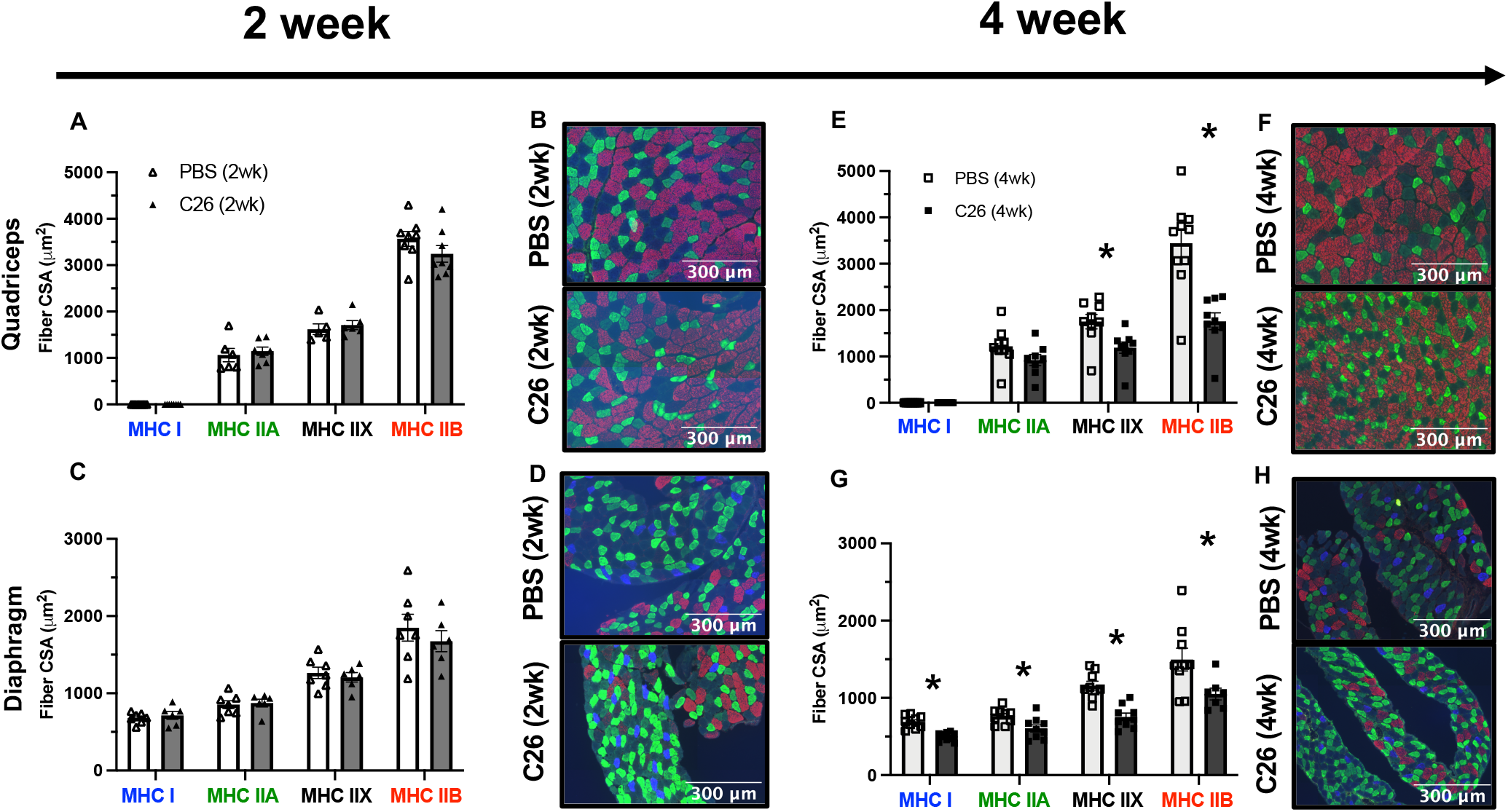
Evaluation of quadriceps and diaphragm fibre-type atrophy in skeletal muscle from C26 tumour-bearing mice. Analysis of fibre histology on MHC isoforms of PBS and C26 mice was performed. Cross-sectional area of MHC stains was evaluated in the quadriceps (*A*, *n*=8; *B*, representative image, magnification × 20) and diaphragm at 2 weeks of tumour bearing (*C*, *n*=6; *D*, representative image, magnification × 20). The same was completed for the quadriceps (*E*, *n*=9; *F*, representative image, magnification × 20) and diaphragm at 4 weeks of tumour bearing (*G*, *n*=9; *H*, representative image, magnification × 20). Results represent mean ± SEM; * *P*<0.05 PBS (4wk) vs C26(4wk).

### Mitochondrial electron transport chain protein contents are reduced in quadriceps but do not change in diaphragm by 4 weeks of tumour development

At 2 weeks, electron transport chain (ETC) subunit contents in both muscles were unchanged in C26 relative to PBS controls (Figure 4A, B). At 4 weeks, C26 showed lower contents in subunits of complex I (−31%), complex II (−18%), complex IV (−37%), complex V (−11%) and total ETC subunit content (−22%; Figure 4C) relative to PBS that were significant or approached significance. ETC subunit contents did not change in diaphragm at 4 weeks relative to PBS (Figure 4D).

**Figure 4.**
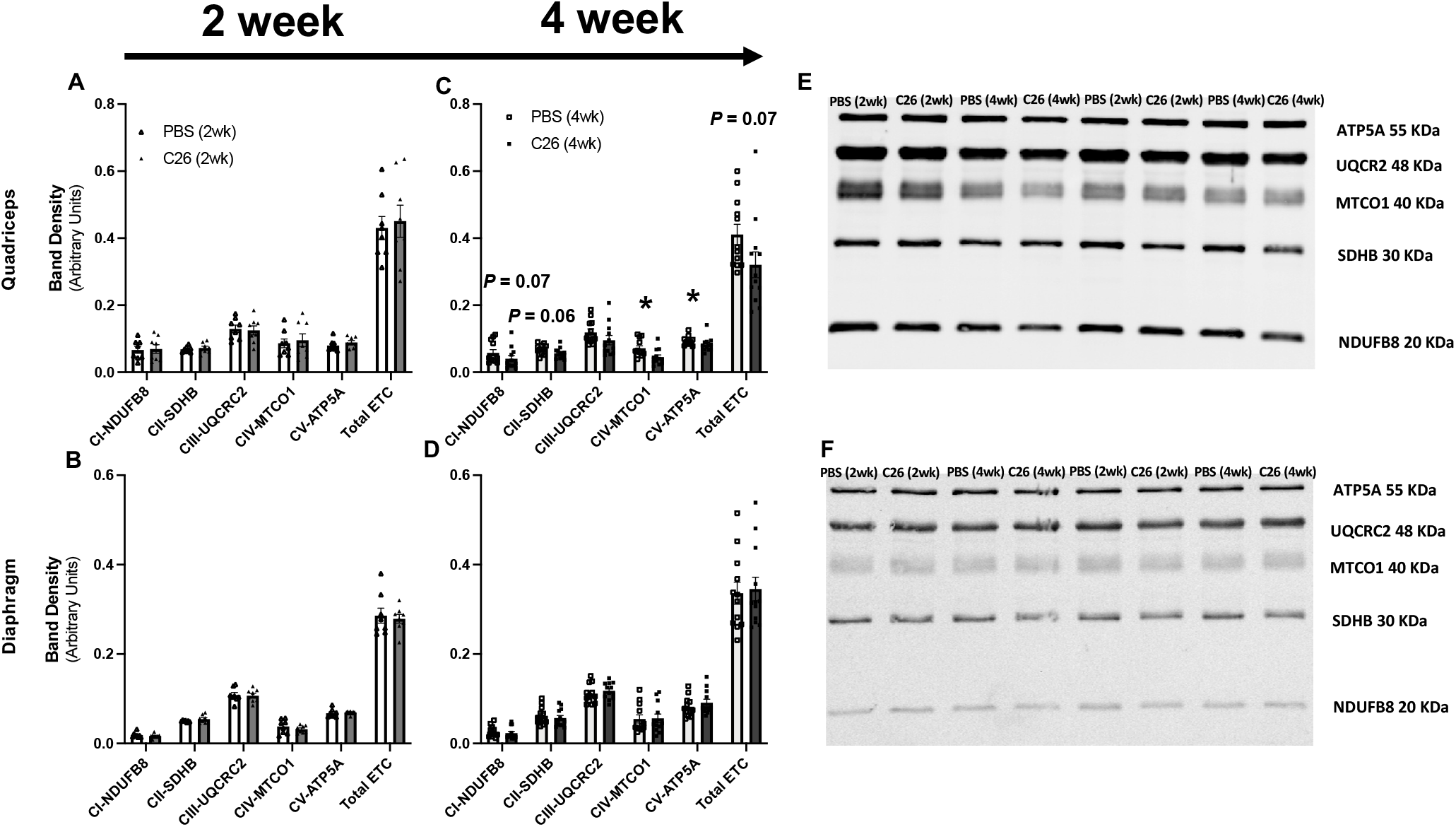
Muscle-specific changes in markers of oxidative phosphorylation in C26 tumour-bearing skeletal muscle. Protein content of electron transport chain components were quantified at in the quadriceps (*A*, *n*=8) and diaphragm at 2 weeks (*B*, *n*=8) and 4 weeks in both muscles respectively *C, D n*=12). *E*, representative image for quadriceps and *F*, representative image for diaphragm. Results represent mean ± SEM; * *P*<0.05 PBS (4wk) vs C26 (4wk).

### Mitochondrial respiratory control by ADP is greater in both muscles by 4 weeks of tumour development despite early reductions in the quadriceps

We determined if the central role of ADP in stimulating respiration was impaired in both quadriceps and diaphragm at 2 and 4 weeks after subcutaneous implantations of C26 cancer cells. We stimulated complex I with NADH generated by pyruvate (5mM) and malate (2mM) across a range of ADP concentrations to challenge mitochondria with a spectrum of metabolic demands. The ADP titrations were repeated with (+Creatine) and without (−Creatine) 20mM creatine to model the two main theoretical mechanisms of energy transfer from mitochondria to cytosolic compartments that utilize or bypass mitochondrial creatine kinase (mtCK) respectively (Figure 5). Briefly, the +Creatine system stimulates mitochondria to export phosphocreatine (PCr) whereas the -Creatine condition drives ATP export.

**Figure 5.**
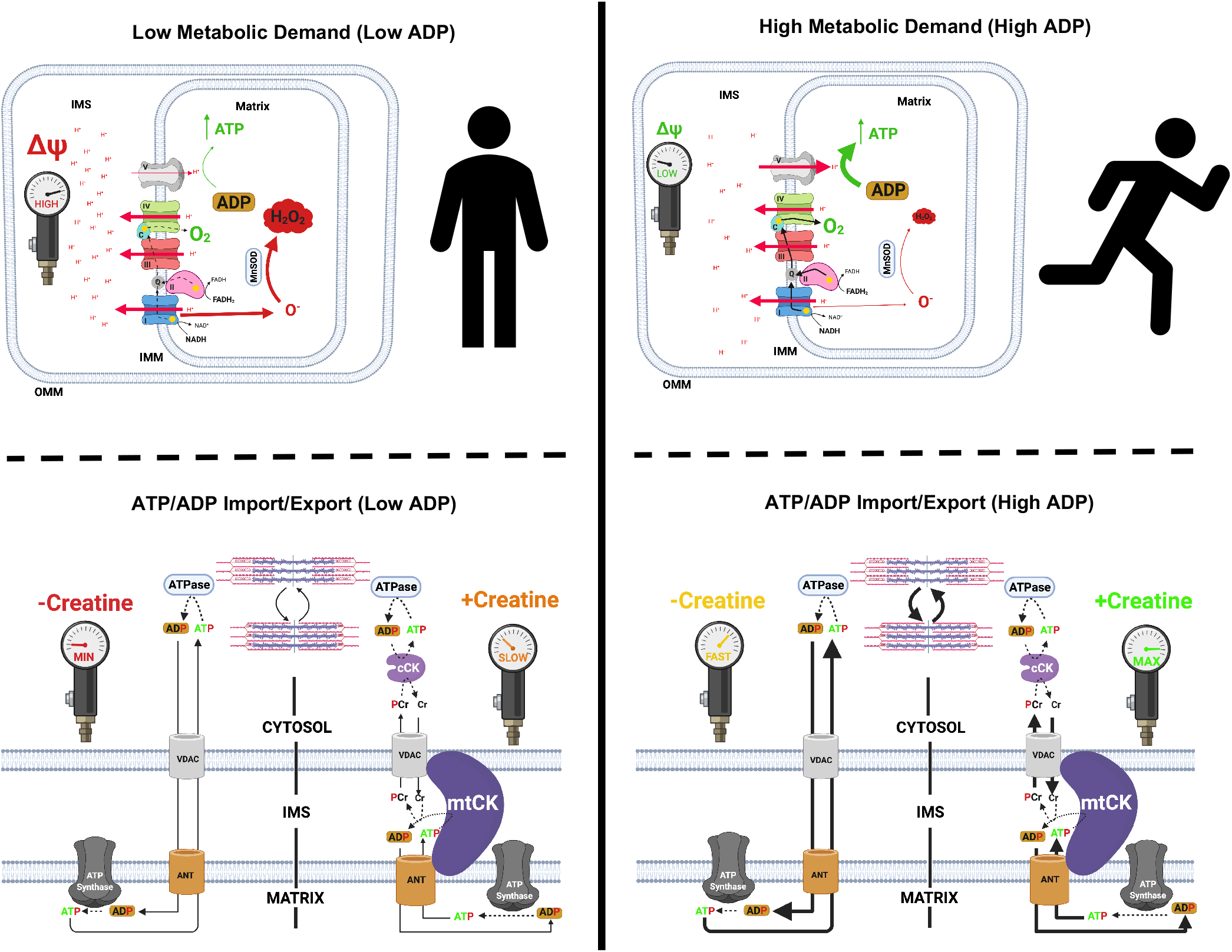
Schematic representation of energy homeostasis in low metabolic (left) vs. high metabolic (right) demand states. When ADP is low, less ATP is produced. A concomitant accumulation of [H^+^] in the inner membrane space (IMS) increases membrane potential (ΔΨ), attenuates H+ pumping, induces premature electron slip and generates superoxide (O^•-^) which is dismutated to H_2_O_2_ by manganese superoxide dismutase (MnSOD; top left). Only Complex I-derived superoxide is displayed. When ADP is high, more ATP is produced as [H^+^] diffuse from the IMS to the mitochondrial matrix through ATP synthase. The decrease in ΔΨ lowers premature electron slip, generating less O^•-^ and H_2_O_2_ (top right). ADP generated by ATPases throughout the cell enter the matrix through the voltage dependent anion channel (VDAC) on the outer mitochondrial membrane (OMM) and the adenine nucleotide translocase (ANT) on the inner mitochondrial membrane (IMM; bottom left). Creatine accelerates matrix ADP/ATP cycling and ATP synthesis by reducing the diffusion distance of the slower diffusing ADP and ATP while shuttling phosphate to the cytoplasm through rapidly diffusing phosphocreatine which is used by cytosolic creatine kinase (cCK) to recycle local ATP to support the activity of various ATPases. Rapidly diffusing creatine returns to the IMS to be re-phosphorylated by mitochondrial creatine kinase (mtCK). Non-ATPase sites of ATP hydrolysis are not displayed but also contribute to net metabolic demand (kinases, and other ATP-dependent processes). The net effect of metabolic demand (global ATP hydrolysis) on matrix ADP/ATP cycling is displayed under the context of creatine independent (−Creatine) and creatine dependent (+Creatine) conditions. Figure adapted from *Aliev et al.*, 2011, *Guzun et al.*, 2012, *Wallimann et al.*, 2011 and Nicholls 2013(23, 43–45). Created with BioRender.com

In both the −Creatine and +Creatine conditions, pyruvate/malate-supported ADP-stimulated respiration normalized per mass of fibre bundles (not corrected for ETC subunit content) was lower 2 weeks after C26 implantations in the quadriceps compared to PBS with a main effect across all ADP concentrations with or without creatine (Figure 6A, B). The general reduction in respiration for C26 normalized per mass of fibre bundle was also seen when data were normalized to ETC subunit content (Figure 6C, D). This suggests respiratory control was reduced within mitochondria due to an inherent property of the ETC not related to ETC abundance. We also evaluated if creatine sensitivity was altered in the C26 tumour-bearing muscle by calculating the +Creatine/-Creatine respiratory ratio. This creatine sensitivity index is a measure of the ability of creatine to stimulate respiration by accelerating matrix ADP/ATP cycling and represents coupling of the creatine kinase system to ATP generation (Figure 5), particularly at sub-maximal ADP concentrations (21, 22) which we have reported previously (17, 18). In quadriceps, creatine sensitivities at 100μM and 500μM ADP were unchanged at 2 weeks in C26 vs PBS at this time point (Figure 6E, F). Collectively, these findings indicate respiration was reduced to similar extents in both −Creatine and +Creatine conditions of energy exchange between mitochondria and cytoplasmic compartments.

**Figure 6.**
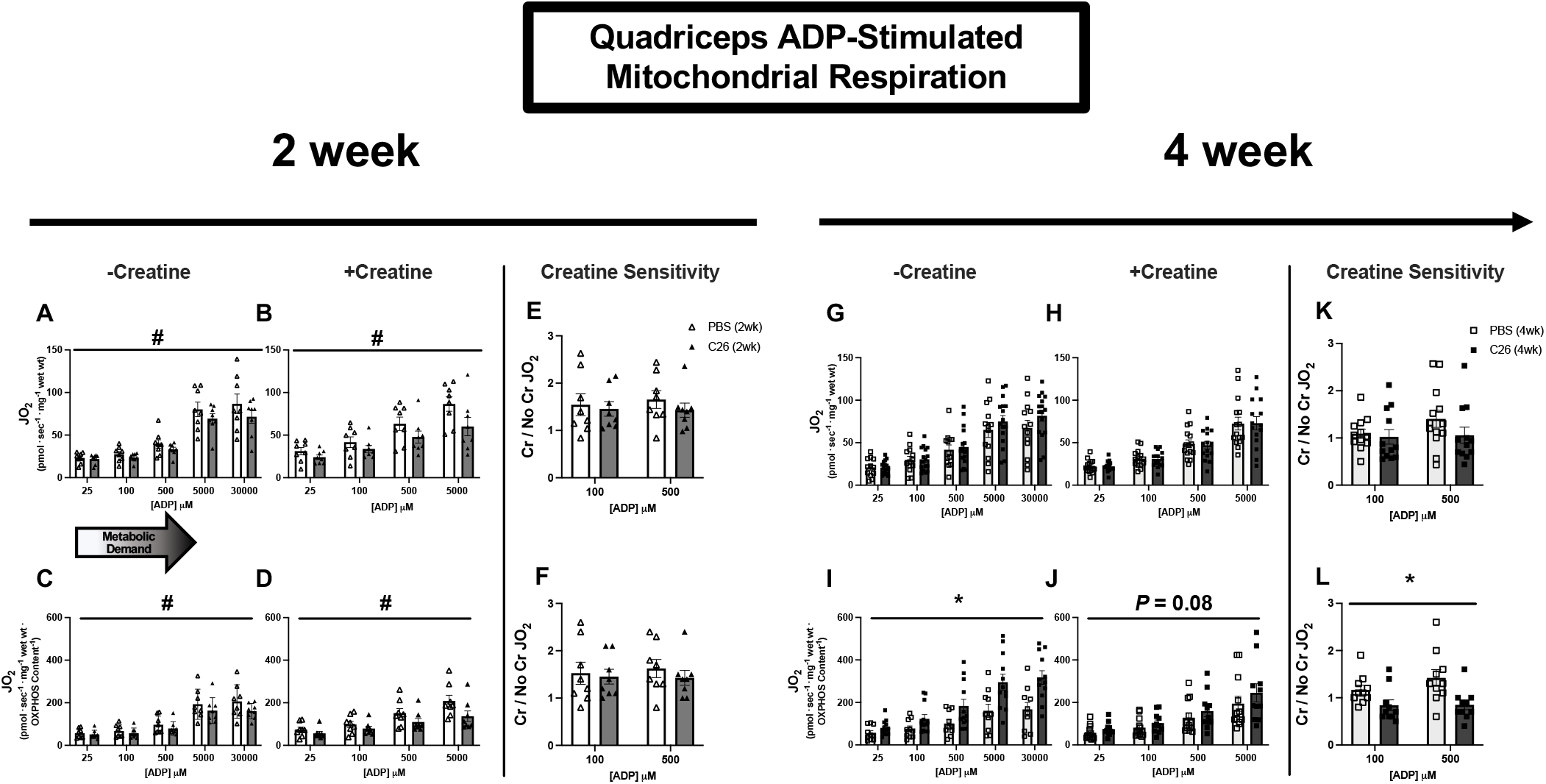
Complex I-supported mitochondrial respiration in quadriceps muscle of C26 tumour-bearing mice. ADP-stimulated (State III) respiration, supported by complex I supported (NADH) substrates pyruvate (5mM) and malate (2mM), was assessed in the absence (−Creatine) and presence (+Creatine) of 20mM creatine at a range of [ADP] until maximal respiration was achieved to model a spectrum of metabolic demands. Respiration was assessed in the quadriceps normalized to bundle size at 2 weeks (*A, B*) and normalized to ETC subunit content (*C, D)* to permit comparisons of intrinsic mitochondrial respiratory responses in each group. Creatine sensitivity was assessed by calculating the +creatine/-creatine ratio (*E, F*) given creatine normally increases ADP-stimulated respiration. The same measurements were completed at 4 weeks (*G-L*). Results represent mean ± SEM; n=8-16; # *P*<0.05, PBS(2wk) vs C26(2wk); * *P*<0.05, PBS(4wk) vs C26(4wk).

These early decrements in quadriceps respiration in C26 at 2 weeks were reversed by 4 weeks. This apparent compensation was seen in both −Creatine and +Creatine conditions. Specifically, respiration was similar to PBS control mice at 4 weeks when normalized per mass of fibre bundle (Figure 6G, H) and higher than controls when normalized to ETC subunit protein content (Figure 6I, J) despite reductions in ETC content as described above (Figure 4C). These findings suggest mitochondria in quadriceps are highly plastic and can super-compensate by upregulating their responsiveness to ADP to levels exceeding PBS controls. Additionally, at 4 weeks, quadriceps mitochondrial creatine sensitivity was impaired in C26 relative to PBS when considering respiration normalized to ETC subunit content given the ratio did not exceed a value of 1.0 which indicates that creatine could not stimulate respiration above the level elicited by ADP alone (Figure 6L). Thus, while C26 cancer strongly increased ADP-stimulated respiration by 4 weeks (Figure 6I), it compromised the coupling of creatine kinase energy transfer, suggesting that this system did not contribute to restored force at this time point (Figure 2A).

In the diaphragm, respiration was similar between C26 and PBS at 2 weeks (Figure 7A-F) but was greater in C26 vs PBS at 4 weeks in both the −Creatine and +Creatine conditions (Figure 7G-J). This upregulation by 4 weeks occurred despite no changes in ETC subunit content as noted above (Figure 4D) which suggests mitochondria increase their responsiveness to ADP through mechanisms that may be independent of mitochondrial content. No changes in creatine sensitivity were observed in C26 vs PBS at 4 weeks (Figure 7K, L) suggesting that coupling of creatine kinase to ATP generation was maintained, in contrast to impaired creatine sensitivity seen in the quadriceps as noted above. Lastly, there was a significant interaction whereby respiration was greater in C26 vs PBS at 5000μM and 7000μM ADP when normalized per mass of fibre bundles (Figure 7G, H) and at all ADP concentrations except 25μM and 100μM when normalized to total ETC subunit content (Figure 7I, J).

**Figure 7.**
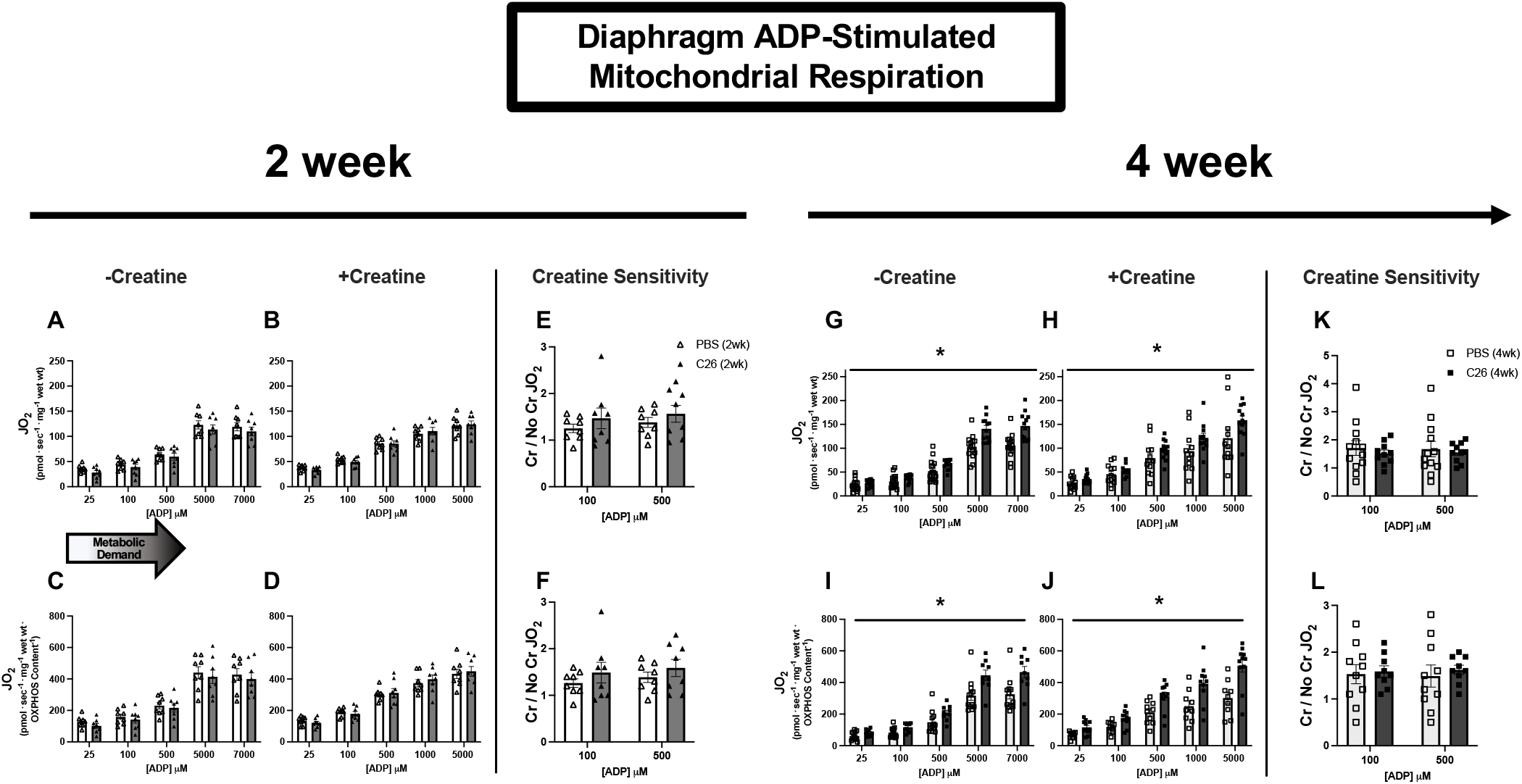
Complex I-supported mitochondrial respiration in diaphragm muscle of C26 tumour-bearing mice. ADP-stimulated (State III) respiration, supported by complex I supported (NADH) substrates pyruvate (5mM) and malate (2mM), was assessed in the absence (−Creatine) and presence (+Creatine) of 20mM creatine at a range of [ADP] until maximal respiration was achieved to model a spectrum of metabolic demands. Respiration was assessed in the diaphragm normalized to bundle size at 2 weeks (*A, B*) and normalized to ETC subunit content (*C, D)* to permit comparisons of intrinsic mitochondrial respiratory responses in each group. Creatine sensitivity was assessed by calculating the +creatine/-creatine ratio (*E, F*) given creatine normally increases ADP-stimulated respiration. The same measurements were completed at 4 weeks (*G-L*). Results represent mean ± SEM; n=8-16; * *P*<0.05, PBS(4wk) vs C26(4wk)

These alterations were specific to pyruvate/malate-supported ADP-stimulated respiration as there were no changes in respiration in response to glutamate (further NADH-generation) and succinate (FADH_2_) generation when comparing C26 to PBS at either time point (Figure S1, S2). By 2 weeks of C26, diaphragm showed a decrease in State II respiration (no ADP, supported by pyruvate/malate; Figure S1, S2) which is generally used as a marker of respiration driven by proton leak into the matrix from the inner membrane space through various sites that are not coupled to ATP synthesis (23). However, State II respiration was greater than control by 4 weeks in both muscles suggesting greater uncoupling at occurs as cancer progresses (Figure S2). Lastly, changes in respiration noted above did not result in changes to the phosphorylation of AMPK in C26 relative to PBS at either 2- or 4-week timepoints (Figure 8); albeit increases in AMPK and the p-AMPK/AMPK ratio were trending in the C26 (4wk) group in the quadriceps.

**Figure 8.**
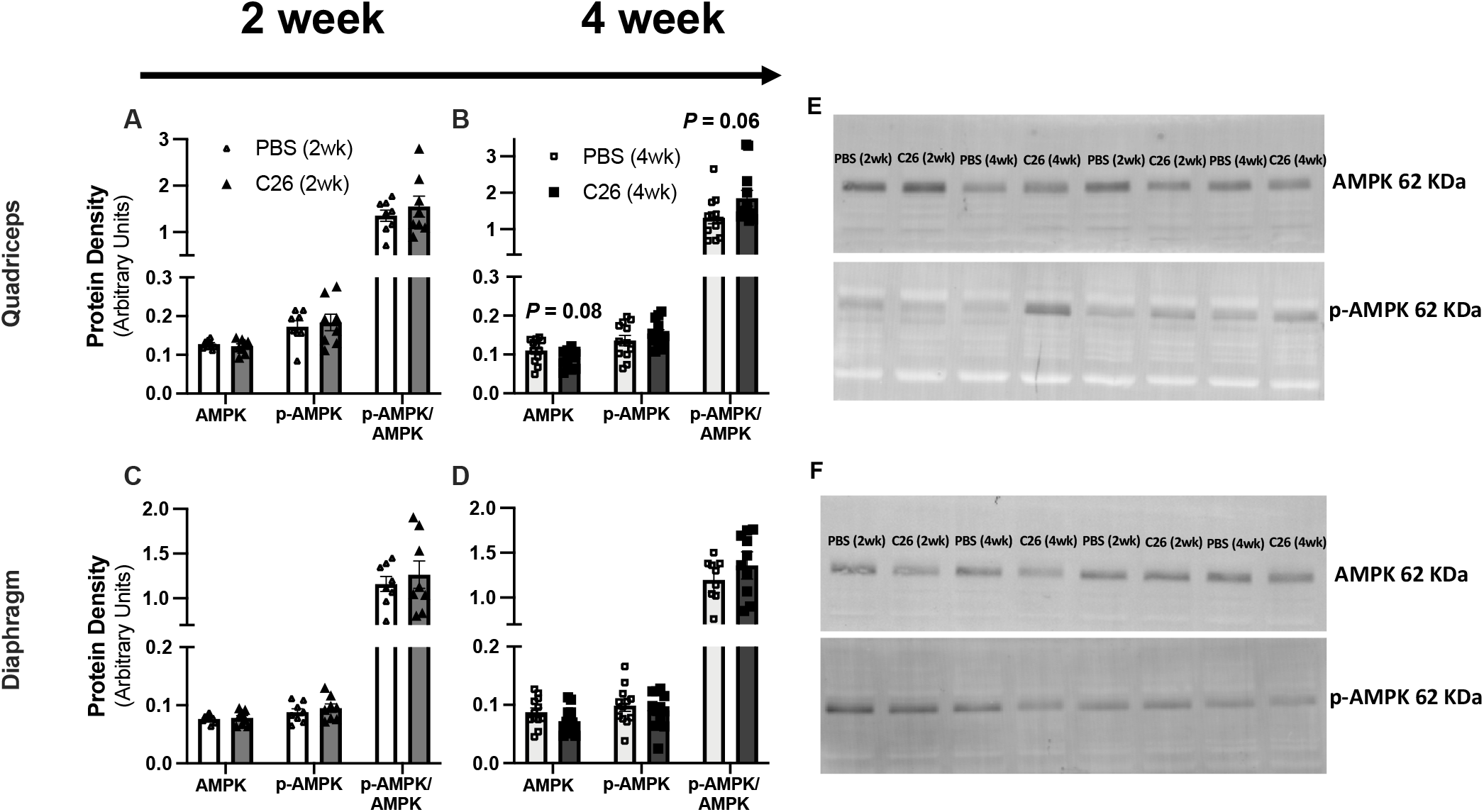
Muscle-specific changes in markers of growth in C26 tumour-bearing skeletal muscle. Protein content of AMPKαand P-AMPKαwere quantified at in the quadriceps at 2 weeks (*A*, *n*=8) and 4 weeks (*B*, *n*=12). Markers were also quantified at in the diaphragm at 2 weeks (*C*, *n*=8) and 4 weeks (*D*, *n*=12). *E*, representative image for quadriceps and *F*, representative image for diaphragm. Results represent mean ± SEM.

### H_2_O_2_ emission is increased in diaphragm early during tumour development and restored to normal by 4 weeks but is unaffected in quadriceps

We stimulated complex I with pyruvate (10mM) and malate (2mM) to generate NADH in the absence of ADP to elicit mH_2_O_2_ emission and determined ADP’s ability to attenuate this emission as occurs naturally during oxidative phosphorylation (see schematic representation, Figure 5). At 2 weeks following C26 implantations, quadriceps mH_2_O_2_ emission was similar to PBS controls under maximal emission conditions (no suppression by ADP, State II; Figure 9A, C) and during suppression by ADP (Figure 9B, D). By 4 weeks of C26 growth, quadriceps mH_2_O_2_ was lower than PBS in both maximal and ADP-suppressive states (Figure 9E, F). However, when mH_2_O_2_ was normalized to total ETC subunit content, no differences were observed between C26 and PBS (Figure 9G, H). This finding suggests eventual decreases in quadriceps mH_2_O_2_ by 4 weeks were related to decreased ETC subunit content as shown in Figur*e* 4. Due to limited tissue availability, pyruvate-supported mH_2_O_2_ was assessed only in the +Creatine condition.

**Figure 9.**
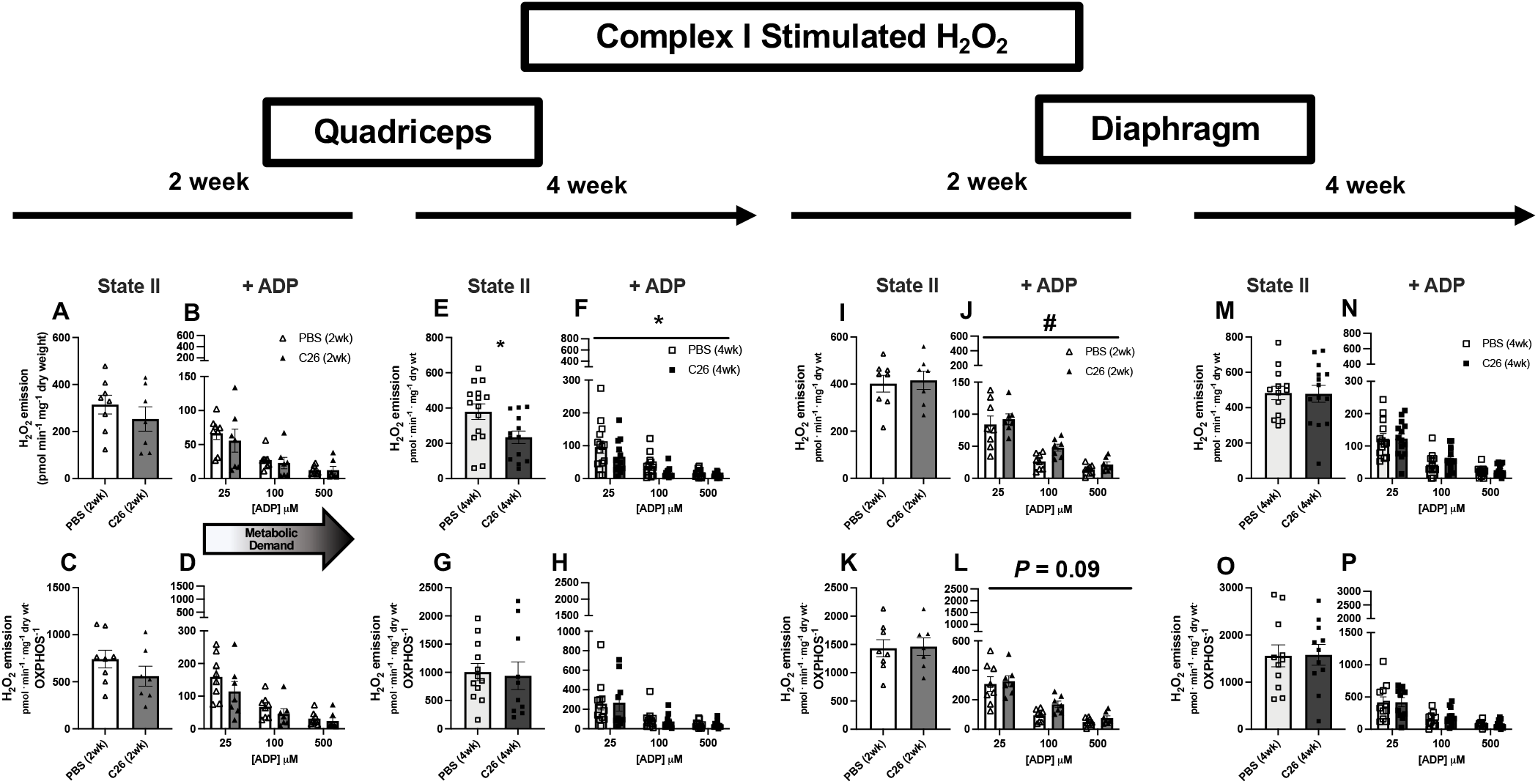
Complex I stimulated mH_2_O_2_ emission in quadriceps and diaphragm muscle of C26 tumour bearing mice. At 2 and 4 weeks, quadriceps mH_2_O_2_ emission supported by pyruvate (10mM) and malate (2mM) (NADH) was assessed under maximal State II (no ADP) conditions in the presence of 20mM creatine (*A, E*) and under a range of [ADP] to model metabolic demand (*B, F*). These measures were also normalized to total ETC subunit content (*C, D, G, H*) to permit comparisons of intrinsic mitochondrial respiratory responses in each group. These measures were repeated in the diaphragm (*I-P*). Results represent mean ± SEM; n=8-16; # *P*<0.05, PBS(2wk) vs C26(2wk); * *P*<0.05, PBS(4wk) vs C26(4wk)

In contrast to the lower mH_2_O_2_ in quadriceps, diaphragm mH_2_O_2_ was greater in C26 mice at 2 weeks relative to PBS in the presence of ADP despite no change in maximal mH_2_O_2_ (Figure 9I, J). This finding reveals C26 causes early elevations in diaphragm mH_2_O_2_ that are likely due to a specific impairment in the ability of ADP to attenuate H_2_O_2_ emission. Moreover, when mH_2_O_2_ emission was normalized to total ETC subunit content at 2 weeks, maximal mH_2_O_2_ emission remained unchanged (Figure 9K), while the higher emissions in the presence of ADP did not reach significance (Figure 9L) but mirrored patterns observed when normalized to wet mass of tissue as noted above. At 4 weeks, there were no differences in diaphragm mH_2_O_2_ under maximal or submaximal (presence of ADP) conditions using either normalization approach (Figure 9M-P) suggesting diaphragm mitochondria are plastic and can eventually restore mH_2_O_2_ to normal levels.

Succinate-supported mH_2_O_2_ emission generally did not change in either muscle in C26 vs PBS at either time point (Figure S3). This finding suggests reverse electron flux to Complex I from Complex II (23) was not altered by C26 cancer, and the responses mentioned above using pyruvate/malate reveal a specific alteration in mH_2_O_2_ emission supported by forward electron flux through Complex I.

## Discussion

Certain indices of skeletal muscle mitochondrial dysfunction have been associated with cancer cachexia in various mouse models (11–13, 24, 25), but the time- and muscle-dependent relationship remains unclear. Here, we demonstrate how quadriceps and diaphragm have both shared and distinct time-dependent responses to cancer in the C26 colon carcinoma mouse model of cancer cachexia. First, weakness was observed prior to atrophy in both muscles, yet an eventual increase in force production to control levels occurred only in quadriceps. Second, atrophy in most fibre types was preceded by altered mitochondrial bioenergetics but the specific relationship differed between muscles with decreases in respiration occurring in quadriceps in contrast to elevated mH_2_O_2_ emission in diaphragm. Third, both muscles upregulated mitochondrial respiration supported specifically by pyruvate and malate substrates at 4 weeks which may reflect a hormetic adaptation to maintain energy homeostasis during cachexia. Likewise, the diaphragm restored Complex I-supported mitochondrial H_2_O_2_ emission to normal lower levels by 4 weeks which demonstrates the transient nature of this potential redox pressure.

Collectively, these findings suggest muscle weakness can occur before atrophy during C26 cancer, and this progression is related to dynamic time-dependent changes in mitochondrial bioenergetics that are unique to each muscle.

### Mitochondrial bioenergetic alterations and skeletal muscle force reductions precede skeletal muscle atrophy

Work from *Brown* et al. suggested mitochondrial degradation and dysfunction precedes muscle atrophy in the LLC xenograft mouse model of cancer cachexia (12). In this study, atrophy markers occurred after earlier indices of mitochondrial degeneration in comparator muscles (flexor digitorum brevis and plantaris) including mitochondrial degradation, respiratory control ratios and H_2_O_2_ emission. The findings of the present study support this proposal with a comparison of atrophy, mitochondrial respiration and mH_2_O_2_ emission within the same muscle types, namely quadriceps and diaphragm. These findings also extend the proposal by showing muscle-specific mitochondrial alterations occur concurrent to muscle weakness and before atrophy. Specifically, early decreases in respiratory kinetics in quadriceps were not seen in diaphragm suggesting that more oxidative muscle might avoid such respiratory decrements. Conversely, early increases in mH_2_O_2_ emission seen in diaphragm did not occur in quadriceps. These relationships suggest targeted therapies to counter mitochondrial alterations during cancer cachexia should consider the specific bioenergetic function that is altered at precise timepoints in each muscle type.

This relationship between early mitochondrial stress prior to atrophy in both muscles becomes further complex when considering force production. Muscle weakness occurred at 2 weeks in both muscles before atrophy which highlights a shared pattern in the progression of muscle dysfunction during cancer. While the purpose of this investigation was not to address other mechanisms regulating force production, reduced fibre sizes cannot be an explanation given atrophy did not occur until after weakness was first observed. However, the distinct mitochondrial signatures in both muscles at 2 weeks could guide additional questions. For example, in the quadriceps, the early reductions in force were associated with early decreases in mitochondrial respiratory control by ADP. When force production was restored to control levels by 4 weeks, respiration actually increased above control levels when normalized to ETC subunit content. This dynamic relationship is intriguing and suggests early quadriceps weakness might be due to impairments in mitochondrial energy provision that is nonetheless plastic and capable of adapting – possibly as a hormetic response to the earlier respiratory deficiency – to correct this weakness through super-compensations in energy supply.

In contrast, the diaphragm weakness seen at 2 weeks might be linked to an early redox pressure given elevated mH_2_O_2_ emission was observed. This observation is consistent with prior observations of early and transient increases in H_2_O_2_ emission in diaphragm in the LLC mouse model of cancer cachexia (24). However, while we did not observe lower respiration in the diaphragm, the increased respiration seen at 4 weeks in this muscle is surprising. The explanation for this increase is not apparent but might suggest an earlier energetic deficiency occurred outside of our selected time points, but this is speculative and would require additional examination.

Overall, while the precise mechanism of lower muscle force is not apparent at 2 weeks in both muscles, the possible contributions of mitochondria could be related more to an early energy crisis in quadriceps vs a redox stress in diaphragm that, as noted above, also preceded the eventual atrophy of each muscle (Figure 10). Additional insight could be gained by extending the current investigation’s focus on fibre type-specific cross sectional area responses to cancer by comparing a wider spectrum of fibre types with regards to mitochondria-atrophy relationships.

**Figure 10.**
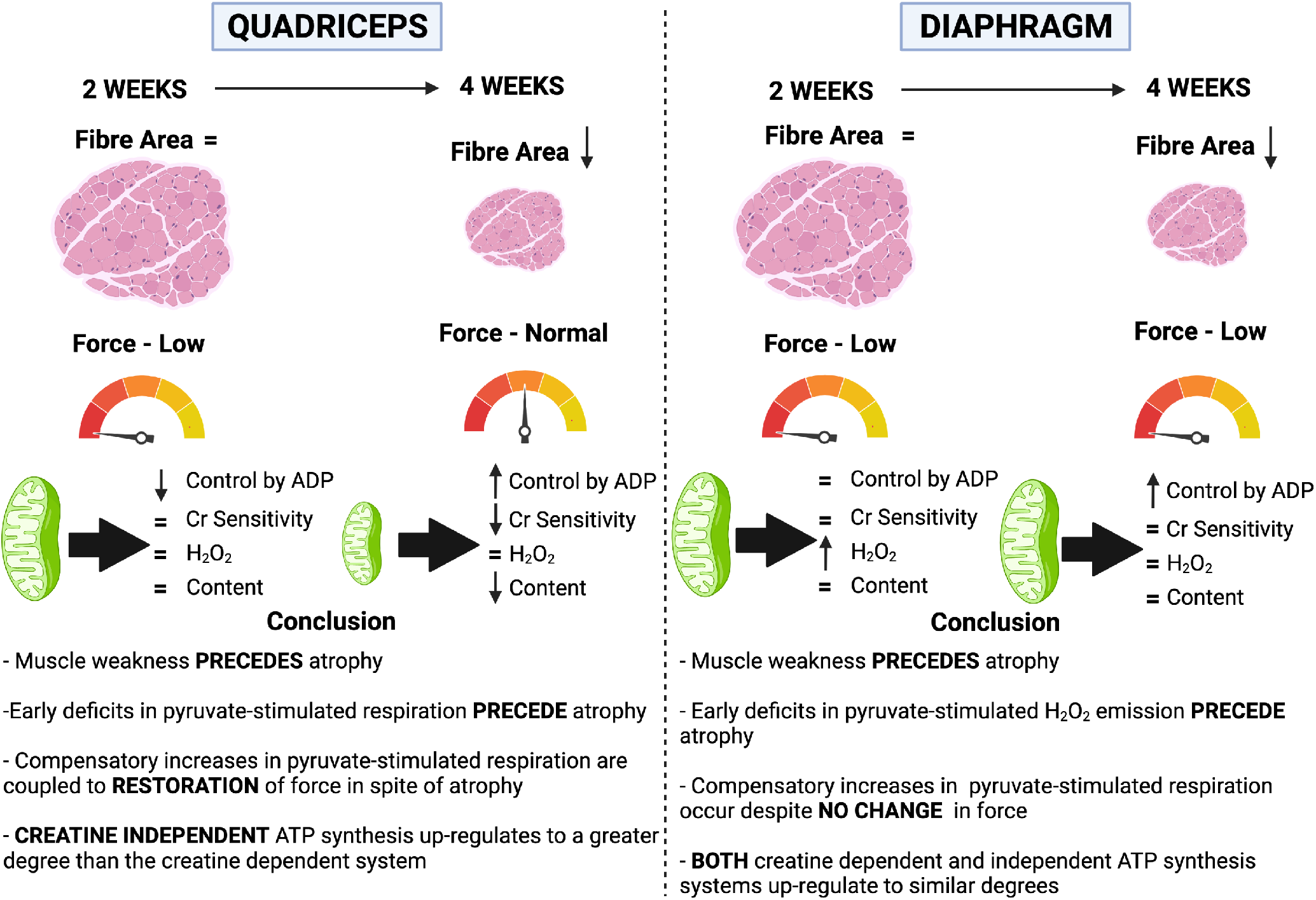
Summary of the time-dependent and muscle-specific adaptations to C26 xenografts in CD2F1 mice. At 2 weeks, early impairments in force generating capacity are associated with reductions in mitochondrial pyruvate/malate-supported ADP-stimulated respiration in quadriceps and elevated mH_2_O_2_ emission in diaphragm. These distinct mitochondrial responses precede atrophy in both muscles by 4 weeks. At this time, quadriceps and diaphragm responses to C26 become heterogeneous. The restoration of force generating capacity in quadriceps in spite of atrophy is not observed in the diaphragm even though both muscles demonstrate apparent compensatory increases in mitochondrial ADP-stimulated respiration. The mitochondrial responses to cancer are more diverse in quadriceps than diaphragm, with increases in respiration by 4 weeks occurring as a potential compensation for reductions in mitochondrial electron transport chain markers. Mitochondrial creatine metabolism is impaired in quadriceps by 4 weeks. Created with BioRender.com

### Perspectives on the potential for mitochondrial hormesis in quadriceps and diaphragm

The findings of lower respiration and increased mH_2_O_2_ emission at 2 weeks is consistent with prior reports at various time points and muscle in the LLC, C26 and peritoneal carcinosis mouse models (11–13, 24, 25). To our knowledge, the eventual increase in pyruvate-supported respiration seen in both quadriceps and diaphragm in the present study is novel, while, the attenuation of mH_2_O_2_ emission seen in the diaphragm is consistent with past reports in the plantaris (12) and diaphragm (24) in the LLC mouse model of cachexia. As noted above, mitochondrial respiratory control by ADP increased above control in both muscles despite a stress being observed earlier only in quadriceps. We questioned whether this early reduction in respiration represented an energy crisis triggering compensatory signaling through the energy sensor AMPK – a pathway that triggers compensatory mitochondrial biogenesis or upregulation of substrate-specific oxidation (26). We did not observe an effect of cancer on AMPK phosphorylation at either time point (Figure 8), although there was a trend in the quadriceps at 4 weeks of tumour bearing whereby AMPK content was increased. Nonetheless, these results do not rule out the potential for AMPK activation at other time points. There are also multiple feedback control systems linking metabolic stress to respiratory control that are independent of AMPK which might be considered in future investigations (27).

The design of substrate titration protocols lends insight into the specific mechanisms by which respiration and mH_2_O_2_ become altered during cancer. For example, as pyruvate/malate was used as the substrates to generate NADH to stimulate complex I-supported respiration, future investigations might consider the potential for cancer to upregulate pyruvate dehydrogenase activity, albeit maximal activity given saturating pyruvate concentrations were used. Also, the consistent increase in respiration across a wide spectrum of ADP concentrations by 4 weeks in both muscles suggests mitochondrial responsiveness to a wide range of metabolic demands may have been enhanced such that key regulators of matrix ADP/ATP cycling could be considered for future directions (Figure 5). ADP was also more effective at attenuating mH_2_O_2_ (23) in quadriceps by 4 weeks (Figure 9) which supports this possibility. Collectively, these findings suggest cancer disrupts mitochondrial bioenergetics by specifically desensitizing mitochondria to ADP in both muscles.

Mitochondrial creatine metabolism appeared to be less capable of adapting in quadriceps by 4 weeks (Figure 6L) suggesting mitochondrial creatine kinase-dependent phosphate shuttling is more affected in this muscle than diaphragm which showed no such deficiency. In fact, the creatine-independent (−Creatine) system showed homogeneous plasticity by upregulating in both muscles by 4 weeks while the creatine-dependent system upregulated only in the diaphragm. These findings suggest mitochondrial creatine metabolism may be disrupted in quadriceps muscle during cancer which may impact energy homeostasis given the importance of this system in certain muscles (21).

In general, the diaphragm appeared to be superior to quadriceps with respect to maintaining mitochondrial ETC content markers and respiratory control by ADP at 2 weeks with evidence of super-compensation in respiratory function at 4 weeks. Furthermore, reductions in ETC protein contents were observed in quadriceps at 4 weeks after C26 implantation whereas no changes were observed in diaphragm. This resilience of diaphragm appears to be a unique finding given prior reports have also shown lower mitochondrial protein markers from various pathways and muscle types in LLC and *APC*^(Min/+)^ mouse models of cancer cachexia (10). While the mechanisms for this muscle heterogeneity remain unclear, one possibility relates to muscle contractile activity. Diaphragm constantly contracts *in vivo* whereas quadriceps is used only during locomotion. As mitochondrial content and substrate oxidation are regulated by contractile activity (27), future directions might consider whether the diaphragm holds a special mitochondrial ‘resistance’ to cancer with respect to energy homeostasis which might support the growing notion of chronic contractile activity in improving muscle health during cancer (28).

In conclusion, this investigation reports muscle weakness precedes atrophy of quadriceps and diaphragm in the C26 colon carcinoma mouse model of cancer cachexia. This progression was associated with heterogenous muscle-specific and time-dependent mitochondrial responses in both muscles. Specifically, an early energetic stress (impaired respiratory control by ADP) was more apparent in quadriceps in contrast to a mitochondrial redox pressure in diaphragm. These early mitochondrial stressors were seemingly corrected as cancer progressed despite the development of atrophy in both muscles and a unique increase in force production in quadriceps. Moreover, C26 cancer caused a unique impairment in the coupling of the mitochondrial creatine kinase system to ATP generation in quadriceps whereas this system was not affected in diaphragm. This dynamic plasticity across time demonstrates how the effects of cancer on one muscle may not predict the response in another muscle type. The findings also highlight how understanding heterogeneity may identify mechanisms that determine whether a given muscle might be sensitive, or resistant, to cancer cachexia.

## Methods

### Animal Care

48 eight-week-old male CD2F1 mice were purchased from Charles River (Massachusetts, USA). Upon arrival, mice were housed and given a minimum of 72 h to acclimatize before cancer implantations. All mice were provided access to standard chow and water ad libitum as differences in food intake has been shown to not impact the C26 model of cancer cachexia (29). Mice were monitored daily for general well-being, tumour ulcerations and tumour size. If mice demonstrated signs of extreme distress, mice would be sacrificed as soon as possible, however, this was never required.

### C26 Cell Culture and Tumour Implantation

C26 cancer cells (Purchased from NCI – Frederick, MD USA) were plated at passage 2-3 in T-75 flasks in DMEM supplemented with 10% foetal bovine serum plus 1% penicillin and streptomycin. Once confluent, cells were trypsinized, counted and diluted in PBS. C26 cells (5 × 10^5^) suspended in 100 μL sterile PBS were implanted subcutaneously to both flanks of mice at 8 weeks of age (11). For control, mice received identical subcutaneous injections of 100 μL sterile PBS and aged for 2 weeks **(PBS (2wk); n= 8)** and 4 weeks (PBS (4wk); n= 16). Tumours developed for 14-17 days (C26 (2wk); n= 8) and 26-29 days (C26 (4wk); n= 16). Tumours were measured daily, recording the length and width of tumours with digital calipers using the following formula to obtain tumour volume (volume of a sphere): (4/3*π*(length/2)*(width/2)2) in accordance with York University Animal Care Committee guidelines. The same investigator was responsible for measuring tumour sizes throughout the length of the study as preliminary work demonstrated that tumour size measurements can vary between individuals (data not shown; CV – 7.2% between 3 individuals, CV – 1.3 within designated individual).

### Surgery Procedure

Quadriceps, soleus, plantaris, gastrocnemius, tibialis anterior, extensor digitorum longus and spleen were quickly collected under isoflurane anesthesia prior to euthanasia. Tissues were weighed and snap-frozen in liquid nitrogen and stored at −80°C. Quadriceps and diaphragm muscles were placed in BIOPS containing (in mM) 50 MES Hydrate, 7.23 K_2_EGTA, 2.77 CaK_2_EGTA, 20 imidazole, 0.5 dithiothreitol, 20 taurine, 5.77 ATP, 15 PCr, and 6.56 MgCl_2_·6 H_2_O (pH 7.1) to be prepared for mitochondrial bioenergetic assays. Quadriceps from one leg and diaphragm strips were harvested for mitochondrial bioenergetic assays while the quadriceps from the contracted leg and a separate diaphragm strip were used for force measurements. The diaphragm strip used for force measurements was cut within 30 seconds of the entire diaphragm being placed in BIOPS prior to transferring the strip to Ringer’s solution as noted below.

### *In Situ* Quadriceps Force and *In Vitro* Diaphragm Force

*In situ* force production for quadriceps muscle was partially adapted from previous literature (30). Mice were anesthetized with isoflurane and shaved of all hair on their hindlimb. An incision was made above the patella to expose the femoral tendon which was then tightly secured with suture. Once the knot was in place, the tendon was carefully severed, and the suture was attached to an Aurora Scientific 305C muscle lever arm with a hook (Aurora, Ontario, Canada). The knee was secured with a vertical knee clamp immobilizing the knee joint with a 27G needle. Contraction of the quadriceps was controlled through percutaneous stimulation of the femoral nerve anterior to the hip joint. Optimal resting length (L_o_) was determined using single twitches (pulse width = 0.2ms) at varying muscle lengths. Once L_o_ was established, force as a function of stimulation frequency was measured during 8 isometric contractions at varying stimulation frequencies (1, 20, 40, 60, 80, 100, 120, 140 Hz). The quadriceps muscle was then weighed and used for normalization of force production.

*In vitro* force production for diaphragm muscle was partially adapted from previous literature (31). Briefly, the diaphragm strip used for force production was placed in a petri dish of ~25°C Ringer’s solution containing (in mM): 121 NaCl, 5 KCl, 1.8 CaCl_2_, 0.5 MgCl_2_ 0,4 NaHPO_4_. 24 NaHCO_3_, 5.5 glucose and 0.1 EDTA; pH 7.3 oxygenated with 95% O_2_ and 5% CO_2_. Diaphragm strips were cut from the central region of the lateral costal hemidiaphragm. Silk suture was tied to the central tendon as well the ribs, and the preparation was transferred to an oxygenated bath filled with Ringer solution, maintained at 25°C. The suture secured to the central tendon was then attached to a lever arm while the suture loop secured to the ribs was attached to a force transducer. The diaphragm strip was situated between flanking platinum electrodes driven by a biphasic stimulator (Model 305C; Aurora Scientific, Inc., Aurora, ON, Canada). Optimal L_o_ was determined using twitches (pulse width = 0.2ms) at varying muscle lengths. Once L_o_ was established, force as a function of stimulation frequency was measured during 10 isometric contractions at varying stimulation frequencies (1, 10, 20, 40, 60, 80, 100, 120, 140, 200 Hz). Force production was normalized to the calculated cross-sectional area (CSA) of the muscle strip (m/l*d) where m is the muscle mass, l is the length, and d is mammalian skeletal muscle density (1.06mg.mm^3^).

### Mitochondrial Bioenergetic Assessments

#### Preparation of Permeabilized Muscle Fibres

The assessment of mitochondrial bioenergetics was performed as described previously in our publications (17, 19, 32). Briefly, the quadriceps and diaphragm from the mouse was removed and placed in BIOPS. Muscle was trimmed of connective tissue and fat and divided into small muscle bundles (~1.2 – 3.7 mg wet weight for quadricep and 0.6 – 2.1 mg for diaphragm). Each bundle was gently separated along the longitudinal axis to form bundles that were treated with 40 μg/mL saponin in BIOPS on a rotor for 30 min at 4°C. Following permeabilization, the permeabilized muscle fibre bundles (PmFB) for respiration assessments were blotted and weighed in ~ 1.5mL of tared pre-chilled BIOPS (muscle relaxing media) to ensure PmFB remained relaxed and hydrated rather than exposed to open air. Wet weights were used given small pieces of muscle can detach during respirometry assessments, albeit greatly reduced by blebbistatin (described below). Mean ± SEM wet weights (mg) were 2.4 ± 0.07 for quadriceps and 1.3 ± 0.04 for diaphragm. The remaining PmFB for mH_2_O_2_ were not weighed at this step as this data was normalized to fully recovered dry weights taken after the experiments. All PmFB were then washed in MiRO5 on a rotator for 15 minutes at 4°C to remove the cytoplasm. MiRO5 contained (in mM) 0.5 EGTA, 10 KH_2_PO_4_, 3 MgCl_2_•6H_2_O, 60 K-lactobionate, 20 Hepes, 20 Taurine, 110 sucrose, and 1 mg/ml fatty acid free BSA (pH 7.1).

#### Mitochondrial Respiration

High-resolution O_2_ consumption measurements were conducted in 2 mL of respiration medium (MiRO5) using the Oroboros Oxygraph-2k (Oroboros Instruments, Corp., Innsbruck, Austria) with stirring at 750 rpm at 37°C. MiRO5 contained 20 mM Cr to saturate mitochondrial creatine kinase (mtCK) and promote phosphate shuttling through mtCK or was kept void of Cr to prevent the activation of mtCK (33) as described in Figure 5. For ADP-stimulated respiratory kinetics, our previously published procedures to stimulate complexes I and II-supported respiration were employed (17–19). 5 mM pyruvate and 2 mM malate were added as complex I-specific substrates (via generation of NADH to saturate electron entry into complex I) followed by a titration of sub-maximal ADP (25, 100 and 500 μM) and maximal ADP (up to 5000 μM in the presence of Cr or 30000 μM in the absence of Cr). 25 μM and 100 μM are close to low and high points of previous estimates of free ADP concentrations in human skeletal muscle in resting and high intensity exercise states and therefore allow the determination of mitochondrial responsiveness to a physiological spectrum of low to high energy demands (34–38). Saturating [ADP] were different depending on the muscle and presence or absence of creatine in the experimental media. Mitochondrial respiration was normalized to mass of fibre bundles as well as total ETC subunit contents to evaluate whether changes in respiration per mass were due to alterations in mitochondrial content or intrinsic mitochondrial respiratory responses.

Kmapp for creatine to ADP was not established as we have observed that many permeabilized fibers from past studies do not fit Michaelis-Menten kinetics with these assay conditions (low to modest R^2^). Creatine accelerates matrix ADP/ATP cycling at submaximal [ADP] and lowers the Kmapp for ADP in some muscles (21, 33). Therefore, in order to evaluate mitochondrial creatine sensitivity, 100 and 500 μM ADP were used to calculate a creatine sensitivity index. Following the ADP titration, cytochrome *c* was added to test for mitochondrial membrane integrity. Finally, succinate (20 mM) was then added to saturate electron entry into Complex II.

All experiments were conducted in the presence of 5 μM blebbistatin (BLEB) in the assay media to prevent spontaneous contraction of PmFB, which has been shown to occur in response to ADP at 37°C that alters respiration rates (33, 39). Polarographic oxygen measurements were acquired in 2 second intervals with the rate of respiration derived from 40 data points and expressed as pmol/s/mg wet weight. PmFB were weighed in ~1.5 mL of tared BIOPS to relax muscle as noted above.

#### Mitochondrial H_2_O_2_ Emission (mH_2_O_2_)

mH_2_O_2_ was determined spectrofluorometrically (QuantaMaster 40, HORIBA Scientific, Edison, NJ, USA) in a quartz cuvette with continuous stirring at 37°C, in 1 mL of Buffer Z supplemented with 10 μM Amplex Ultra Red, 0.5 U/ml horseradish peroxidase, 1mM EGTA, 40 U/ml Cu/Zn-SOD1, 5 μM BLEB and 20mM Cr. Buffer Z contained (in mM) 105 K-MES, 30 KCl, 10 KH_2_PO_4_, 5 MgCl_2_ • 6H_2_O, 1 EGTA, and 5mg/mL BSA (pH 7.4). State II mH_2_O_2_ (maximal emission in the absence of ADP) was induced using the Complex I-supporting substrates (NADH) pyruvate (10mM) and malate (2mM) to assess maximal (State II, no ADP) mH_2_O_2_ as described previously (18). Following the induction of State II mH_2_O_2_, a titration of ADP was employed to progressively attenuate mH_2_O_2_ as occurs when membrane potential declines during oxidative phosphorylation (Figure 5). After the experiments, the fibers were rinsed in double deionized H_2_O, lyophilized in a freeze-dryer (Labconco, Kansas City, MO, USA) for > 4h and weighed on a microbalance (Sartorius Cubis Microbalance, Gottingen Germany). The rate of mH_2_O_2_ emission was calculated from the slope (F/min) using a standard curve established with the same reaction conditions and normalized to fibre bundle dry weight.

### Western Blotting

A frozen piece of quadriceps and diaphragm from each animal was homogenized in a plastic microcentrifuge tube with a tapered Teflon pestle in ice-cold buffer containing (mm) 20 Tris/HCl, 150 NaCl, 1 EDTA, 1 EGTA, 2.5 Na_4_O_7_P_2_, 1 Na_3_VO_4_, 1% Triton X-100 and PhosSTOP inhibitor tablet (Millipore Sigma, Burlington, MA, USA) (pH 7.0) as published previously (40). Protein concentrations were determined using a bicinchoninic acid (BCA) assay (Life Technologies, Carlsbad, CA, USA). 15-30 μg of denatured and reduced protein was subjected to 10-12% gradient SDS-PAGE followed by transfer to low-fluorescence polyvinylidene difluoride membrane. Membranes were blocked with Odyssey Blocking Buffer (LI-COR, Lincoln NE, USA) and immunoblotted overnight (4°C) with antibodies specific to each protein. A commercially available monoclonal antibody was used to detect electron transport chain proteins (rodent OXPHOS Cocktail, ab110413; Abcam, Cambridge, UK, 1:250 dilution), including V-ATP5A (55kDa), III-UQCRC2 (48kDa), IV-MTCO1 (40kDa), II-SDHB (30 kDa), and I-NDUFB8 (20 kDa). Commercially available polyclonal antibodies were used to detect AMP-activated protein kinaseα(AMPKα) (rabbit, CST, 2532; 62kDa; 1:1000) and Phospho-AMPKαThr172 (P-AMPK) (rabbit CST, 2535, 62kDa; 1:500) as used previously (40).

After overnight incubation in primary antibodies, membranes were washed 3×5 minutes in TBST and incubated for 1 hour at room temperature with the corresponding infrared fluorescent secondary antibody (LI-COR IRDye 680nm or 800nm) at a dilution previously optimized (1:20 000). Immunoreactive proteins were detected by infrared imaging (LI-COR CLx; LI-COR) and quantified by densitometry using ImageJ. All images were normalized to Amido Black total protein stain (A8181, Sigma) using the entire lane corresponding to each sample.

### Immunofluorescence Analysis

Quadriceps (vastus intermedius & vastus lateralis) and diaphragm muscle samples embedded in O.C.T medium (Fisher Scientific) were cut into 10-μm- thick sections with a cryostat (HM525 NX, Thermo Fisher Scientific, Mississauga, ON, Canada) maintained at −20°C. Muscle fibre type was determined as previously described (41), with minor modifications. All primary antibodies were purchased from the Developmental Studies Hybridoma Bank (University of Iowa), and secondary antibodies were purchased from Invitrogen (Burlington, ON, Canada). Briefly, slides were blocked with 5% goat serum (Sigma Aldrich) in PBS for 1 hour at room temperature. Next, slides were incubated with primary antibodies against myosin heavy chain (MHC) I (BA-F8; 1:25), MHC IIA (SC-71; 1:1000) and MHC IIB (BF-F3; 1:50) for 2 hours at room temperature. Afterwards, slides were washed 3x in PBS for 5 minutes and then incubated with secondary antibodies (MHCI; Alexa Fluor 350 IgG2b; 1:1000), (MHCIIa; Alexa Fluor 488 IgG1; 1:1000), (MHC Iib; Alexa Fluor 568 IgM; 1:1000) for 1 hour at room temperature. Slides were then washed 3x in PBS for 5 minutes and mounted with ProLong antifade reagent (Life Technologies, Burlington, ON, Canada). Images were acquired the next day using EVOS FL Auto 2 Imaging System (Invitrogen, Thermo Fisher Scientific, Mississauga, ON, Canada). Individual images were taken across the entire cross section and then assembled into a composite image. 20-30 muscle fibers per fiber type were selected randomly throughout the cross section and traced with ImageJ software to assess CSA after calibrations with a corresponding scale bar. Muscle fibers that appeared black were recorded as MHC IIX (41).

### Statistics

Results are expressed as mean ± SEM. The level of significance was established at *P* < 0.05 for all statistics. The D’Agostino – Pearson omnibus normality test was first performed to determine whether data resembled a Gaussian distribution. Western blot results for proteins in the electron transport chain subunit complexes I, IV and V in quadriceps failed normality as did proteins in complexes I, II, IV and V for diaphragm. In addition, quadriceps and diaphragm delta glutamate respiration failed normality and were analyzed using a non-parametric Mann-Whitney t-test. All other data passed normality. A two-tailed unpaired t-test was used to compare C26 to PBS within each time point with respect to muscle mass, fibre cross-sectional area, and remaining western blots. Mitochondrial respiration, mH_2_O_2_ and force-frequency were analyzed using a two-way ANOVA with factors of timepoint (2 vs 4 week) and treatment (C26 vs PBS) followed by Benjamini, Krieger and Yekutieli’s post-hoc analysis when a main affect was obtained to identify a significant interaction between groups (GraphPad Prism Software 8.4.2, La Jolla, CA, USA) (42).

### Study Approval

All experiments and procedures were approved by the Animal Care Committee at York University (AUP Approval Number 2019-10) in accordance with the Canadian Council on Animal Care.

## Author Contributions

L.J.D., C.A.B, M.R.C., N.P.G. and C.G.R.P. contributed to the rationale and study design. L.J.D., C.A.B., S.G. and C.G.R.P. conducted all experiments and/or analyzed all data. C.G.R.P. and L.J.D. wrote the manuscript. All authors contributed to the interpretation of the data and manuscript preparation. All authors have approved the final version of the manuscript and agree to be accountable for all aspects of the work. All persons designated as authors qualify for authorship, and all those who qualify for authorship are listed.

## Acknowledgments

Funding was provided to C.G.R.P. by the National Science and Engineering Research Council (no. 436138-2013 and no.2019-06687) and an Ontario Early Researcher Award (C.G.R.P., no. 2017-0351) with infrastructure supported by the Canada Foundation for Innovation, the Ontario Research Fund and the James. H. Cummings Foundation. L.J.D. was supported by a NSERC CGS-M scholarship. C.A.B. was supported by a NSERC PGS-D scholarship. S.G. was supported by an Ontario Graduate Scholarship. N.P.G. was funded by U.S. National Institutes of Health under Award Number R01AR075794 from the National Institute of Arthritis and Musculoskeletal and Skin Diseases.

## Supplemental Figure Legends

**Figure S1** Multiple substrate evaluation of oxygen consumption in quadriceps permeabilized muscle fibre bundles. Oxygen consumption was evaluated in the absence of creatine at 2 weeks and 4 weeks post C26 implantation or PBS injections in permeabilized muscle fibres when stimulated with glutamate (*A, B*), succinate (*E, F*) and pyruvate/malate (*I, J*). This was repeated in the presence of 20mM Creatine (*C, D, G, H, K, L*). Results represent mean ± SEM; n=8-16; # *P*<0.05 PBS (2wk) vs C26 (2wk); * *P*<0.05 PBS (4wk) vs C26 (4wk)

**Figure S2** Multiple substrate evaluation of oxygen consumption in diaphragm permeabilized muscle fibre bundles. Oxygen consumption was evaluated in the absence of creatine at 2 weeks and 4 weeks post C26 implantation or PBS injections in permeabilized muscle fibres when stimulated with glutamate (*A, B*), succinate (*E, F*) and pyruvate/malate (*I, J*). This was repeated in the presence of 20mM Creatine (*C, D, G, H, K, L*). Results represent mean ± SEM; n=8-16; # *P*<0.05 PBS (2wk) vs C26 (2wk); * *P*<0.05 PBS (4wk) vs C26 (4wk).

**Figure S3** Succinate stimulated mH_2_O_2_ emission in quadriceps and diaphragm muscle of C26 tumour bearing mice. At 2 and 4 weeks, quadriceps mH_2_O_2_ emission supported by succinate (10mM) (FADH_2_) was assessed under maximal State II (no ADP) conditions in the absence of creatine (*A, E*) and in the presence of 20mM Creatine (*C, G*). State III (range of [ADP] to model metabolic demand) was also assessed in the absence of creatine (*B, F*) and in the presence of 20mM creatine (*D* and *H*). These measures were repeated in the diaphragm (I-P). Results represent mean ± SEM; n=7-16; # *P*<0.05, PBS (2wk) vs C26 (2wk); * *P*<0.05, PBS (4wk) vs C26 (4wk).

## Notes

### Competing Interest Statement

The authors have declared no competing interest.

## References

1. Fearon K, et al. Definition and classification of cancer cachexia: An international consensus. Lancet Oncol. 2011;12(5):489–495. doi:10.1016/S1470-2045(10)70218-7

2. Kasvis P, et al. Health-related quality of life across cancer cachexia stages. Ann Palliat Med. 2019;8(1):33–42. doi:10.21037/apm.2018.08.04

3. Dewys WD, et al. Prognostic effect of weight loss prior to chemotherapy in cancer patients. Eastern Cooperative Oncology Group. Am J Med. 1980;69(4):491–497. doi:10.1016/s0149-2918(05)80001-3

4. Baracos VE, et al. Cancer cachexia is defined by an ongoing loss of skeletal muscle mass. Ann Palliat Med Vol 8, No 1 (January 2019) Ann Palliat Med (Update Cancer Cachexia Mem Ken Fearon). Published online 2018. https://apm.amegroups.com/article/view/22915

5. Arthur ST, et al. Cachexia among US cancer patients. J Med Econ. 2016;19(9):874–880. doi:10.1080/13696998.2016.1181640

6. Tan BHL, Fearon KCH. Cachexia: Prevalence and impact in medicine. Curr Opin Clin Nutr Metab Care. 2008;11(4):400–407. doi:10.1097/MCO.0b013e328300ecc1

7. Tisdale MJ. Molecular pathways leading to cancer cachexia. Physiology. 2005;20:340–348. doi:10.1152/physiol.00019.2005

8. Argilés JM, et al. Cancer cachexia: Understanding the molecular basis. Nat Rev Cancer. Published online 2014:754–762. doi:10.1038/nrc3829

9. Halle JL, et al. Tissue-specific dysregulation of mitochondrial respiratory capacity and coupling control in colon-26 tumor-induced cachexia. Am J Physiol Regul Integr Comp Physiol. 2019;317(1):R68–R82. doi:10.1152/ajpregu.00028.2019

10. White JP, et al. Muscle oxidative capacity during IL-6-dependent cancer cachexia. Am J Physiol - Regul Integr Comp Physiol. 2011;300(2):R201–R211. doi:10.1152/ajpregu.00300.2010

11. Neyroud D, et al. Colon 26 adenocarcinoma (C26)-induced cancer cachexia impairs skeletal muscle mitochondrial function and content. J Muscle Res Cell Motil. 2019;40(1):59–65. doi:10.1007/s10974-019-09510-4

12. Brown JL, et al. Mitochondrial degeneration precedes the development of muscle atrophy in progression of cancer cachexia in tumour-bearing mice. J Cachexia Sarcopenia Muscle. 2017;8(6):926–938. doi:10.1002/jcsm.12232

13. Smuder AJ, et al. Pharmacological targeting of mitochondrial function and reactive oxygen species production prevents colon 26 cancer-induced cardiorespiratory muscle weakness. Oncotarget. 2020;11(38):3502–3514. doi:10.18632/oncotarget.27748

14. Ballarò R, et al. Targeting Mitochondria by SS-31 Ameliorates the Whole Body Energy Status in Cancer- and Chemotherapy-Induced Cachexia. Cancers (Basel). 2021;13(4):850. doi:10.3390/cancers13040850

15. Toyama EQ, et al. AMP-activated protein kinase mediates mitochondrial fission in response to energy stress. Science. 2016;351(6270):275–281. doi:10.1126/science.aab4138

16. Hancock CR, et al. High-fat diets cause insulin resistance despite an increase in muscle mitochondria. Proc Natl Acad Sci. 2008;105(22):7815 LP - 7820. doi:10.1073/pnas.0802057105

17. Hughes MC, et al. Impairments in left ventricular mitochondrial bioenergetics precede overt cardiac dysfunction and remodelling in Duchenne muscular dystrophy. J Physiol. 2020;598(7):1377–1392. doi:10.1113/JP277306

18. Hughes MC, et al. Early myopathy in Duchenne muscular dystrophy is associated with elevated mitochondrial H2O2 emission during impaired oxidative phosphorylation. J Cachexia Sarcopenia Muscle. 2019;10(3):643–661. doi:10.1002/jcsm.12405

19. Ramos S V, et al. Mitochondrial bioenergetic dysfunction in the D2.mdx model of Duchenne muscular dystrophy is associated with microtubule disorganization in skeletal muscle. PLoS One. 2020;15(10):e0237138. doi:10.1371/journal.pone.0237138

20. Turnbull PC, et al. The fatty acid derivative palmitoylcarnitine abrogates colorectal cancer cell survival by depleting glutathione. Am J Physiol Cell Physiol. 2019;317(6):C1278–C1288. doi:10.1152/ajpcell.00319.2019

21. Kuznetsov A V, et al. Striking differences between the kinetics of regulation of respiration by ADP in slow-twitch and fast-twitch muscles in vivo. Eur J Biochem. 1996;241(3):909–915. doi:10.1111/j.1432-1033.1996.00909.x

22. Schlattner U, et al. Mitochondrial Proteolipid Complexes of Creatine Kinase. Subcell Biochem. 2018;87:365–408. doi:10.1007/978-981-10-7757-9_13

23. Nicholls DG, Ferguson SJ. Bioenergetics (Fourth Edition).; 2013.

24. Rosa-Caldwell ME, et al. Mitochondrial Function and Protein Turnover in the Diaphragm are Altered in LLC Tumor Model of Cancer Cachexia. Int J Mol Sci. 2020;21(21):7841. doi:10.3390/ijms21217841

25. Julienne CM, et al. Cancer cachexia is associated with a decrease in skeletal muscle mitochondrial oxidative capacities without alteration of ATP production efficiency. J Cachexia Sarcopenia Muscle. 2012;3(4):265–275. doi:10.1007/s13539-012-0071-9

26. Steinberg GR, Kemp BE. AMPK in Health and Disease. Physiol Rev. 2009;89(3):1025–1078. doi:10.1152/physrev.00011.2008

27. Hargreaves M, Spriet LL. Skeletal muscle energy metabolism during exercise. Nat Metab. 2020;2(9):817–828. doi:10.1038/s42255-020-0251-4

28. Mina DS, et al. Exercise as part of routine cancer care. Lancet Oncol. 2018;19(9):e433–e436. doi:10.1016/S1470-2045(18)30599-0

29. Talbert EE, et al. Modeling human cancer cachexia in colon 26 tumor-bearing adult mice. J Cachexia Sarcopenia Muscle. 2014;5(4):321–328. doi:10.1007/s13539-014-0141-2

30. Pratt SJP, Lovering RM. A stepwise procedure to test contractility and susceptibility to injury for the rodent quadriceps muscle. J Biol methods. 2014;1(2):e8. doi:10.14440/jbm.2014.34

31. Fajardo VA, et al. Diaphragm assessment in mice overexpressing phospholamban in slow-twitch type I muscle fibers. Brain Behav. 2016;6(6):e00470. doi:10.1002/brb3.470

32. Ramos S V, et al. Altered skeletal muscle microtubule-mitochondrial VDAC2 binding is related to bioenergetic impairments after paclitaxel but not vinblastine chemotherapies. Am J Physiol - Cell Physiol. 2019;316(3):C449–C445. doi:10.1152/ajpcell.00384.2018

33. Perry CGR, et al. Inhibiting myosin-ATPase reveals a dynamic range of mitochondrial respiratory control in skeletal muscle. Biochem J. 2011;437(2):215–222. doi:10.1042/BJ20110366

34. Walsh B, et al. The role of phosphorylcreatine and creatine in the regulation of mitochondrial respiration in human skeletal muscle. J Physiol. 2001;537(3):971–978. doi:https://doi.org/10.1111/j.1469-7793.2001.00971.x

35. Perry CGR, et al. High-intensity aerobic interval training increases fat and carbohydrate metabolic capacities in human skeletal muscle. Appl Physiol Nutr Metab = Physiol Appl Nutr Metab. 2008;33(6):1112–1123. doi:10.1139/H08-097

36. Chen Z-P, et al. Effect of Exercise Intensity on Skeletal Muscle AMPK Signaling in Humans. Diabetes. 2003;52(9):2205 LP – 2212. doi:10.2337/diabetes.52.9.2205

37. Jubrias SA, et al. Acidosis inhibits oxidative phosphorylation in contracting human skeletal muscle in vivo. J Physiol. 2003;553(2):589–599. doi:https://doi.org/10.1113/jphysiol.2003.045872

38. Wackerhage H, et al. Recovery of free ADP, Pi, and free energy of ATP hydrolysis in human skeletal muscle. J Appl Physiol. 1998;85(6):2140–2145. doi:10.1152/jappl.1998.85.6.2140

39. Hughes MC, et al. Mitochondrial Bioenergetics and Fiber Type Assessments in Microbiopsy vs. Bergstrom Percutaneous Sampling of Human Skeletal Muscle. Front Physiol. 2015;6:360. doi:10.3389/fphys.2015.00360

40. Houde VP, et al. AMPK β1 reduces tumor progression and improves survival in p53 null mice. Mol Oncol. 2017;11(9):1143–1155. doi:10.1002/1878-0261.12079

41. Bloemberg D, Quadrilatero J. Rapid determination of myosin heavy chain expression in rat, mouse, and human skeletal muscle using multicolor immunofluorescence analysis. PLoS One. 2012;7(4):e35273. doi:10.1371/journal.pone.0035273

42. Glickman ME, et al. False discovery rate control is a recommended alternative to Bonferroni-type adjustments in health studies. J Clin Epidemiol. 2014;67(8):850–857. doi:https://doi.org/10.1016/j.jclinepi.2014.03.012

43. Aliev M, et al. Molecular system bioenergics of the heart: experimental studies of metabolic compartmentation and energy fluxes versus computer modeling. Int J Mol Sci. 2011;12(12):9296–9331. doi:10.3390/ijms12129296

44. Guzun R, et al. Regulation of respiration in muscle cells in vivo by VDAC through interaction with the cytoskeleton and MtCK within Mitochondrial Interactosome. Biochim Biophys Acta - Biomembr. 2012;1818(6):1545–1554. doi:https://doi.org/10.1016/j.bbamem.2011.12.034

45. Wallimann T, et al. The creatine kinase system and pleiotropic effects of creatine. Amino Acids. 2011;40(5):1271–1296. doi:10.1007/s00726-011-0877-3

